# Differential effects of acute and prolonged morphine withdrawal on motivational and goal-directed control over reward-seeking behavior

**DOI:** 10.1101/2023.09.14.557822

**Authors:** Briac Halbout, Collin Hutson, Stuti Agrawal, Zachary A. Springs, Sean B. Ostlund

**Affiliations:** Department of Anesthesiology and Perioperative Care, School of Medicine, University of California, Irvine, Irvine, CA, 92697, USA; Department of Neurobiology and Behavior, School of Biological Sciences, University of California, Irvine, Irvine, CA, 92697, USA

## Abstract

Opioid addiction is a relapsing disorder marked by uncontrolled drug use and reduced interest in normally rewarding activities. The current study investigated the impact of spontaneous withdrawal from chronic morphine exposure on emotional, motivational, and cognitive processes involved in regulating the pursuit and consumption of natural food rewards in male rats. In Experiment 1, rats experiencing acute morphine withdrawal lost weight and displayed somatic signs of drug dependence. However, hedonically-driven sucrose consumption was significantly elevated, suggesting intact and potentially heightened emotional reward processing. In Experiment 2, rats undergoing acute morphine withdrawal displayed reduced motivation when performing an effortful response for palatable food reward. Subsequent reward devaluation testing revealed that acute withdrawal also disrupted their ability to exert flexible goal-directed control over their reward-seeking behavior. Specifically, morphine-withdrawn rats displayed insensitivity to reward devaluation both when relying on prior action-outcome learning and when given direct feedback about the consequences of their actions. In Experiment 3, rats tested after prolonged morphine withdrawal displayed heightened rather than diminished motivation for food rewards and retained their ability to engage in flexible goal-directed action selection. However, brief re-exposure to morphine was sufficient to impair motivation and disrupt goal-directed action selection, though in this case insensitivity to reward devaluation was only observed in the presence of morphine-paired context cues and in the absence of response-contingent feedback. We suggest that these opioid-withdrawal induced deficits in motivation and goal-directed control may contribute to addiction by interfering with the pursuit of adaptive alternatives to drug use.

## INTRODUCTION

Opioid addiction is a major public health crisis with devastating costs for individuals, communities, and society at large [1,2]. A defining characteristic of opioid addiction is *goal-narrowing*, which involves the excessive and uncontrolled urge to use drugs [3] as well as reduced interest in alternative, non-drug rewards [4-6]. This loss of adaptive goal-directed behavior is multifaceted, comprising distinct emotional, motivational, and cognitive components [7,8], and is positively correlated with withdrawal symptoms and craving in abstinent opioid users [9,10]. Cognitive impairments associated with opioid use tend to persist into drug abstinence and are often exacerbated during early withdrawal [11-15]. However, even after prolonged drug abstinence, exposure to opioid-related cues can trigger cognitive dysfunction [16] and induce a disruptive attentional bias that is associated with eventual relapse [17,18].

Preclinical research indicates that repeated opioid exposure can cause persistent behavioral and cognitive aberrations [6,19,20], though it remains unclear precisely how such treatments impact the way adaptive rewards are valued and pursued. For instance, animals with a history of opioid exposure display either diminished [21-24] or heightened [25-30] palatable food reward seeking and consumption, depending on study parameters such as withdrawal interval. Specifically, feeding and food motivation tend to be depressed during the first few days of opioid withdrawal, but recover and may even become elevated after more prolonged withdrawal [31-34]. Opioid withdrawal may also impact certain feeding processes differently than others. For instance, studies focusing on hedonic-emotional measures of feeding have typically found either no change or increased responsivity in animals undergoing opioid withdrawal [29,35], whereas homeostatic feeding tends to be suppressed [31,33].

Repeated opioid exposure can also disrupt action selection [36-42]. However, it remains unclear whether or how chronic opioid exposure specifically impacts the goal-directed processes that support flexible decision making based on expected behavioral outcomes. This cognitive capacity for goal-directed control can be readily assayed using the reward devaluation task [43], which requires animals to evaluate the *current* value of potential outcomes using previously learned action-outcome relationships. It has been proposed that this capacity for adaptive goal-directed control breaks down in drug addiction [4,44-46], such that drug pursuit becomes disconnected from the many adverse consequences of this behavior. Such an impairment may also *indirectly* contribute to goal-narrowing in addiction by disrupting the processes through which alternative non-drug goals are evaluated and flexibly pursued.

The current study examined the impact of spontaneous withdrawal from chronic morphine exposure (10-30 mg/kg) on hedonic, motivational, and cognitive processes supporting the pursuit and consumption of palatable food rewards in male rats. Experiment 1 characterized the behavioral signs of early acute morphine withdrawal (24-48 h) and examined how this treatment impacted hedonic feeding behavior. Experiments 2 and 3 examined motivational vigor and goal-directed control over instrumental food-seeking actions during early (Experiment 2) and late (Experiment 3) stages of morphine withdrawal. The influence of morphine-related contextual cues on goal-directed choice was also characterized. Our findings indicate that motivation and goal-directed action selection are both disrupted during early but not late morphine withdrawal. However, even after prolonged withdrawal, brief re-exposure to morphine was sufficient to reestablish a reduction in motivational vigor as well as a partial, context-dependent deficit in goal-directed choice.

## MATERIALS AND METHODS

Please see Supplementary Information for additional methods.

### Subjects

Adult male (300-440g) Long-Evans rats (N = 65) were used as subjects. Ad lib food and water were provided except when rats were food-restricted for behavioral procedures (see below). All experimental procedures were approved by the UC Irvine Institutional Animal Care and Use Committee and conducted in accordance with the National Research Council Guide for the Care and Use of Laboratory Animals.

### Morphine treatment

During initial morphine exposure (11d; Fig 1A), rats were injected immediately before being placed in the behavioral chamber for 30 min. Rats were treated twice per day. Visual, tactile, and olfactory cues were added to the chambers to create two distinctive contexts (Contexts A and B). Morphine groups (Fig 1B) were injected with saline each morning before being placed in the *unpaired context (Context A or B)*. Each afternoon these rats were injected with morphine a placed in the alternate, *paired context*. Morphine dose increased over days (10-30 mg/kg) [29,32]. Saline groups received saline injections in both the morning (*unpaired context)* and afternoon (*paired context)*.

**Figure 1.**
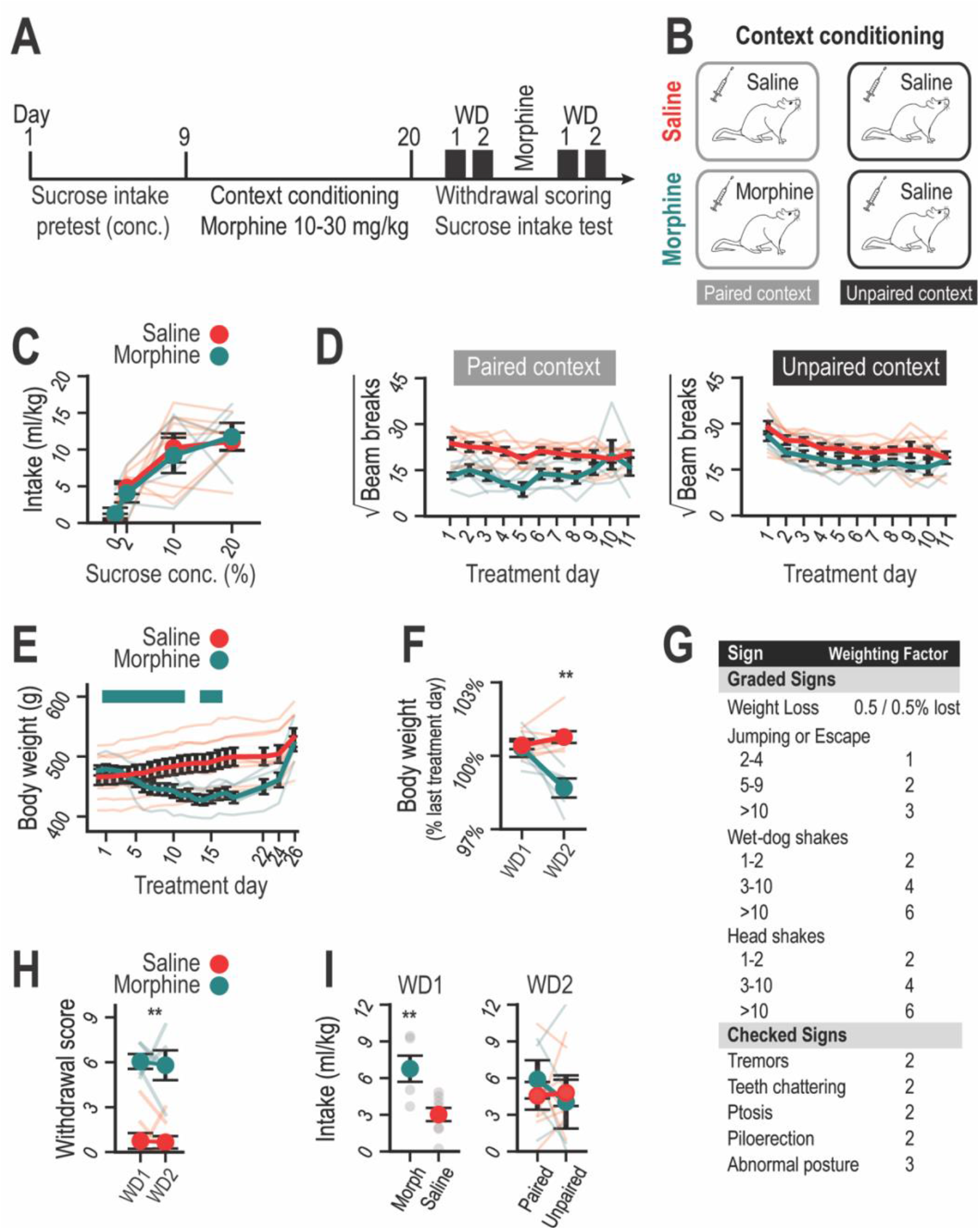
Effects of morphine exposure on somatic withdrawal signs and hedonic feeding in rats. **A**. Schematic representation of behavioral testing during withdrawal from morphine exposure in rats. Rats were tested for somatic signs and sucrose intake during withdrawal days (WDs) 1 and 2. **B**. Schematic diagram illustrating context conditioning in rats from the Morphine or Saline groups. **C**. Initial sucrose intake increases across a range of concentrations (conc.) during pretesting (i.e., before morphine exposure) but does not differ with planned treatment groups (Concentration: F_*3,33*_ = 24.01, p < .001. Concentration x Group and. Group, Fs < 1). **D**. Locomotor activity is differently altered by morphine and saline treatments in paired or unpaired contexts (Day x Context x Group: F_1,110_ = 3.55, p < .001). It is initially reduced by morphine treatment (paired context; Day x Group: F_10,110_ = 3.05, p = .002). However, activity in the unpaired context is similar following morphine and saline treatment (Group: F_1,11_ = 2.26, p = .16; Day: F_10,110_ = 122.43, p < .001; Day x Group: F_10,110_ < 1). **E**. Morphine treatment induces significant weight loss that persists during withdrawal (Group x Day (1-11): F_10,110_ = 47.16, p < .001). Horizontal green mark shows morphine treatment days. **F**. Discontinuation of morphine treatment elicits further weight loss which is apparent on WD2 (% of last treatment day for WD1 and WD2; Group x Day (2): F_1,11_ = 16.41, p = .002). **G**. Graded and Checked withdrawal signs and their corresponding weighing factors for withdrawal scoring. **H**. Morphine treatment induces significant somatic withdrawal signs on WDs 1 and 2 (Group: F_1,11_ = 70.07, p < .001; Day and Day x Group: F’s < 1). **I**. Sucrose intake is significantly elevated in WD1 (t_,11_ =3.30, p = .007), but is unaffected by WD2, regardless of test context (Fs < 1). **p < .01.

### Experiment 1

#### Sucrose intake training

Ad lib fed rats were trained to consume 20% sucrose solution from a retractable metal drinking spout (5 days; 30-min each), before assessing the influence of sucrose concentrations (0, 2, 10, 20%, randomized order). Our primary measure was bodyweight-normalized sucrose intake during the first 3 min of licking behavior, as in [47,48], which selective assays hedonic feeding with minimal influence of satiety [49,50].

#### Morphine exposure

Rats in the morphine group (n = 5) received one saline injection and one morphine injection each day, which were paired with distinct context cues. In contrast, a saline-only group (n = 8) received two saline injections each day, which were also paired with the two distinct contexts.

#### Sucrose intake during withdrawal

Within 24-h of the last injection (WD1), rats underwent morphine withdrawal assessment (see below) and followed by a 2% sucrose intake test. On the next day (WD2), rats underwent the same procedures except that the *paired* or *unpaired* contexts were added to chambers during sucrose intake testing. Rats were then re-exposed to morphine (15 mg/kg for 1d and 30 mg/kg for 2d) and/or saline using the context-treatment arrangements in place during initial drug exposure. Rats were tested in the bare chamber on WD1 and then in the alternate context on WD2, so that each rat was tested in both paired and unpaired contexts.

#### Morphine withdrawal assessment

Prior to each sucrose intake test (WD1 and WD2), rats were placed in a transparent plastic cylinder and continuously video recorded over a 30-min observation period. Trained observers blind to the treatment scored withdrawal signs [51,52] (weighting factors shown in Fig 1G). Withdrawal scores were averaged for each withdrawal interval (WD1 and WD2).

### Experiment 2

#### Instrumental training

Rats were food restricted and underwent instrumental training as in [48,53-55] in bare behavioral chambers (no context cues). After magazine training, rats were trained on two distinct instrumental action-outcome contingencies (e.g., left press → grain pellet and right press → 50% sweetened condensed milk, or vice versal) (Figure 2A and B), using increasingly more effortful random ratio (RR) schedules reward over days (RR5→RR10→RR20).

**Figure 2.**
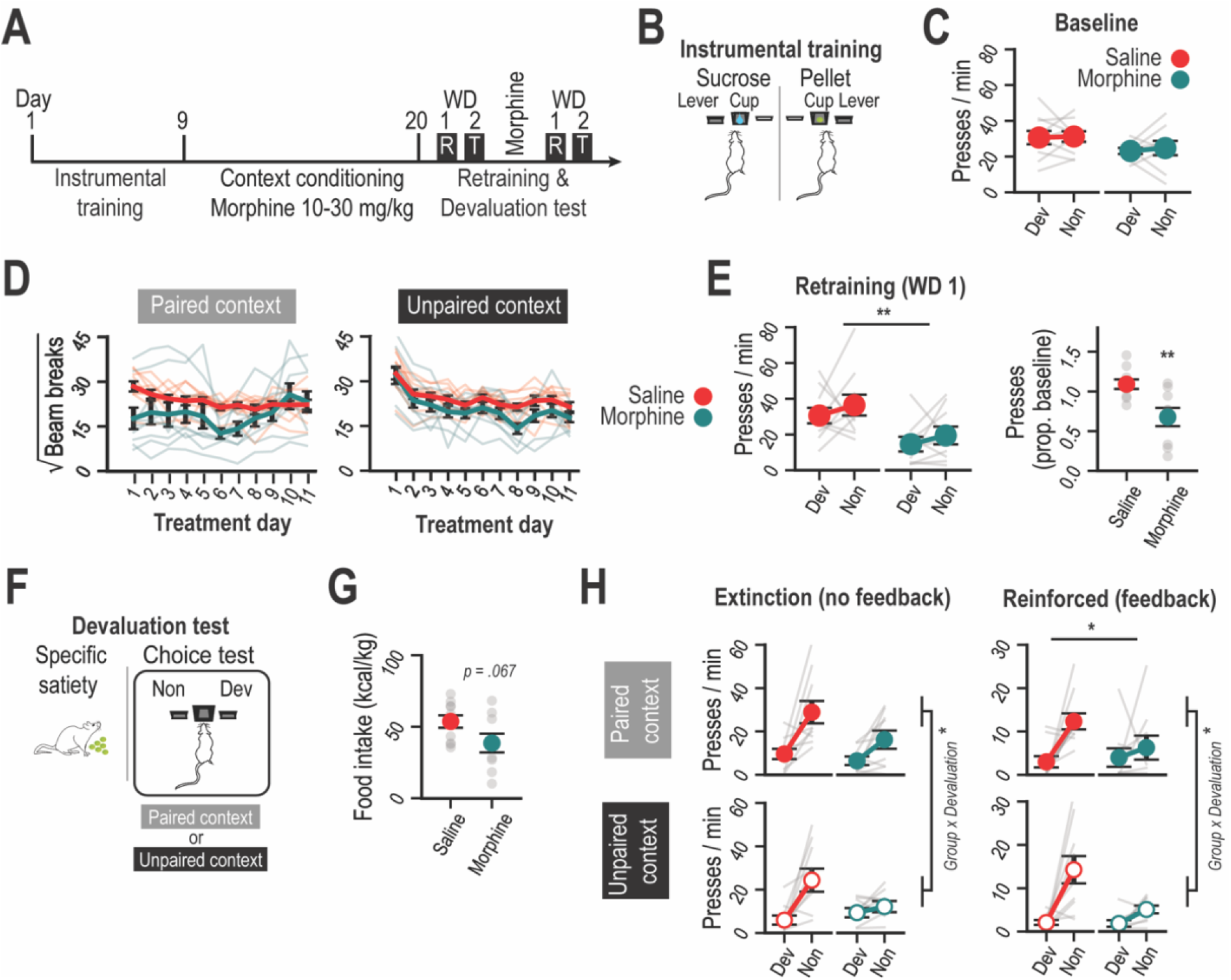
Early morphine withdrawal impairs motivation and goal-directed control over instrumental behavior in rats. **A**. Schematic representation of behavioral testing during early morphine withdrawal. Rats were assessed for motivation vigor during instrumental retraining on withdrawal day (WD) 1. On WD2, rats were given a reward devaluation test in either paired or unpaired contexts to assess their capacity for goal-directed control. **B**. Schematic representation of instrumental training contingencies. **C**. Baseline press rates during initial instrumental training did not differ between groups (Group: F_1,17_ = 3.26, p = .089) or planned devaluation treatment (Devaluation and Devaluation x Group: F’s < 1). **D**. As in Experiment 1, locomotor activity is differently altered during morphine and saline treatments in paired or unpaired contexts (Day x Context x Group: F_10,170_ = 4.58, p < .001). For paired context sessions, morphine treated rats displayed a suppression of activity that was less apparent over days (Day x Group: F_10,170_ = 3.12, p = .001). The groups did not differ and showed similar rates of habituation of locomotor activity during unpaired context sessions (Group: F_1,17_ = 3.19, p = .09; Day: F_10,170_ = 20.75, p < .001; Day x Group: F_10,170_ = 1.024, p < .43). **E**. Instrumental performance is reduced in morphine-treated rats during instrumental retraining on WD1. Left, press rates are separated by group and by planned devaluation conditions, showing an overall decrease in press rate (Group: F_1,17_ = 9.78, p = .006), but not as a function of planned devaluation treatment (Devaluation: F_1,17_ = 1.40, p = .25; Devaluation x Group: F<1). Right, press rates were averaged across actions and plotted as a proportion of baseline performance (t_17_ = 3.32, p = .004). **F**. Schematic of the outcome-specific reward devaluation test. **G**. Food intake during the prefeeding period (specific-satiety induction) was slightly reduced in morphine treated rats (t_17_ = 1.96, p = .067). **H**. Morphine-treated rats show impaired sensitivity to reward devaluation (see main text for statistical analysis). Press rates during the test session are separated by devaluation (Dev or Non), group (Morphine or Saline), test phase (Extinction and Reinforced), and context (Paired or Unpaired). R = Retraining; T = Test; Dev = Devalued; Non = Nondevalued. *p < .05. **p < .01.

#### Morphine exposure

After instrumental training, rats were returned to ad lib food access before receiving 11d of both saline and morphine injections (morphine group: n = 9) or saline-only injections (n = 10), using distinct treatment-context pairing as described above. Rats were returned to food restriction on treatment day 10 and remained restricted for the rest of the experiment.

#### Devaluation tests

On WD1, rats were given instrumental retraining with both action-outcome contingencies. On WD2, rats were given a reward devaluation test as in previous studies [43,48,53,54,56]. Rats were given 60 min unrestricted access to grain pellets or SCM (counterbalanced with drug treatment and training contingencies) to induce outcome-specific satiety. Rats were then placed in the chamber with Context-A cues (either morphine-paired or unpaired, counterbalanced with drug, satiety, and training conditions) (Figure 2F). Both levers were inserted after a 10-min exploration period. For the next 5 min (*Extinction Phase*), rats were able to freely press the left and right lever but received no food reinforcement/feedback. This was immediately followed by a 15-min period (*Reinforced Phase*), during which rats received feedback, as each action was reinforced with its respective outcome. Rats were then re-exposed to morphine and/or saline (with the same context-treatment pairings; as in Experiment 1). They were then retrained on WD1 before undergoing a second round of reward devaluation testing on WD2, this time in the presence of Context-B cues.

### Experiment 3

#### Instrumental training

Rats were food-restricted and given instrumental training as described in Experiment 2. Ad libitum access to food was then provided following the last training sessions.

#### Initial morphine exposure

Rats were given 11 d of saline and morphine injections (morphine group: n = 13) or saline-only injections (n = 13) in distinct context. Rats were rested for the next 14 d before returning to food restriction.

#### Devaluation tests

Rats received instrumental retraining sessions during withdrawal days 19-21, and a devaluation test in Context-A on WD22, as described in Experiment 2 (see Figure 3A). Rats were retrained before undergoing a second devaluation test on WD26 in Context-B.

**Figure 3.**
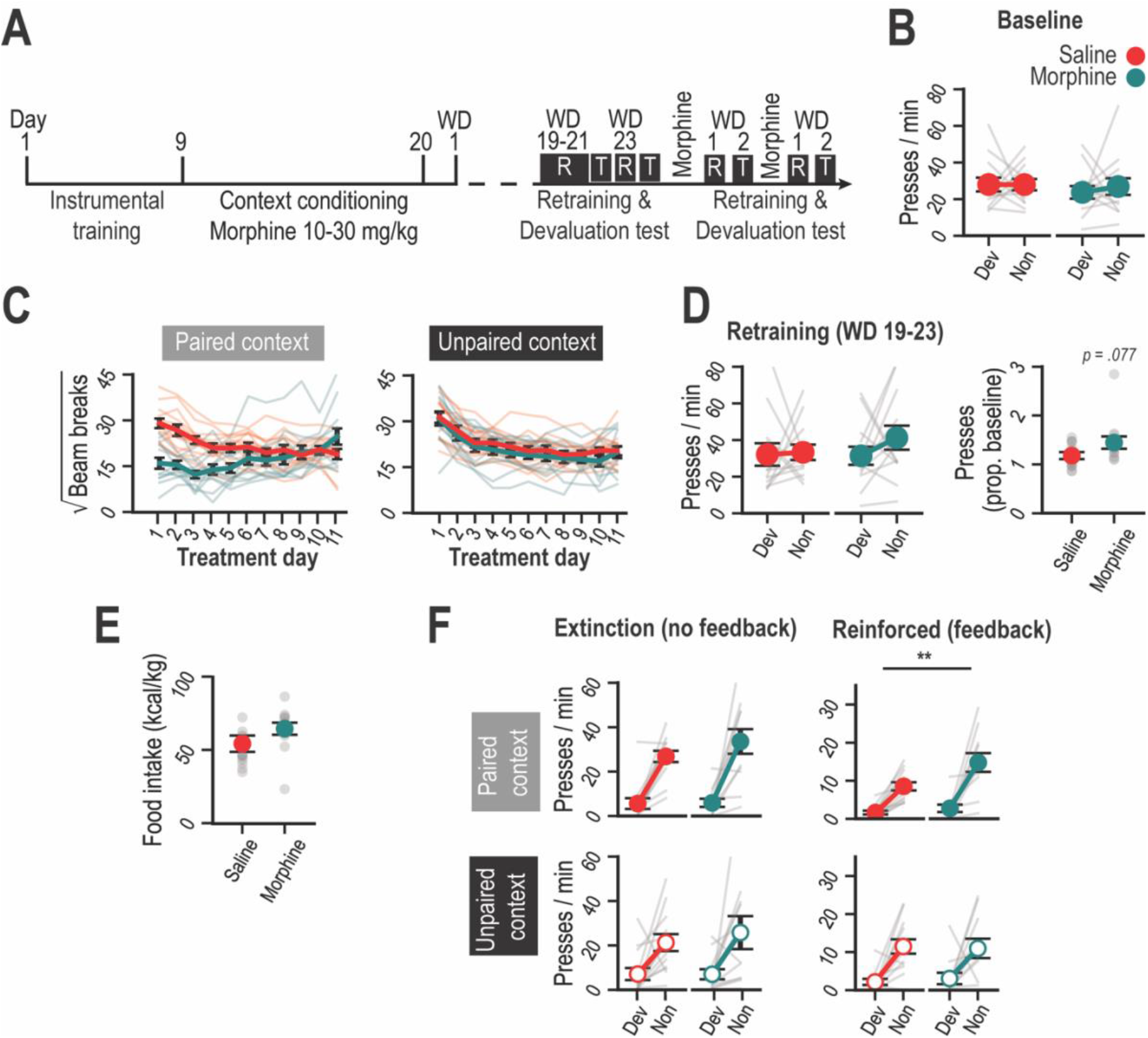
Prolonged morphine withdrawal spares motivation and goal-directed control in rats. **A**. Schematic of behavioral testing during prolonged morphine withdrawal. Rats were assessed for motivation vigor during instrumental retraining on withdrawal days (WDs) 19-23. On WDs 22 and 24, rats were given a reward devaluation test in either paired or unpaired contexts to assess their capacity for goal-directed control. Rats were retested during early withdrawal following morphine re-exposure (see Figure 4). **B**. Baseline press rates during initial instrumental training did not differ between groups (all F’s < 1). **C**. Locomotor activity during morphine and saline treatments in paired or unpaired contexts. Morphine treatment influences locomotor activity, which varies across treatment days (Day x Context x Group: F_10,240_ = 9.91, p < .001). For paired context sessions, morphine injections initially suppress locomotor activity (Day x Group: F_10,240_ = 17.03, p < .001). Saline and morphine groups show a similar decline in activity over days in the unpaired context (Day: F_10,240_ = 43.55, p < .001; Group and Day x Group interactions: F’s < 1). **D**. Instrumental performance during instrumental retraining on WDs 19-21 & 23. Left, press rates in rats separated by group and by planned devaluation conditions do not differ (F<1). Right, press rates averaged across actions and plotted as a proportion of baseline performance show a slight elevation in responding in the morphine group (t_24_=1.85, p = 0.077). **E**. Food intake during the prefeeding period (specific-satiety induction) does not differ between groups (t_24_ = 1.48, p = .15). **F**. Morphine-treated rats show intact sensitivity to reward devaluation (see main text for statistical analysis). Press rates during the test session are separated by devaluation (Dev or Non), group (Morphine or Saline), test phase (Extinction and Reinforced), and context (Paired or Unpaired). R = Retraining; T = Test; Dev = Devalued; Non = Nondevalued. **p < .01.

#### Morphine re-exposure and devaluation tests

To assess the impact of acute withdrawal after brief morphine re-exposure on instrumental performance, rats were given 3 d of treatment with morphine (15 mg/kg for 1d and 30 mg/kg for 2d) and saline (or saline only) using the original treatment-context pairings. Rats then received one day of instrumental retraining on WD1, as in Experiment 2, followed by a devaluation test in Context-A on WD2. Rats were then re-exposed to morphine (30 mg/kg) and saline (or saline only) for 2 d and then retrained on WD1 before undergoing a final devaluation test in Context-B on WD2 (n =13, saline-only group; n = 11, morphine group).

### Data analysis

Data were analyzed using mixed ANOVAs or unpaired t-tests, as appropriate, in SPSS(v29), with significance set at p<.05. Group sizes (n) indicate rats that completed testing and were included in final analysis. See Supplementary Information for details.

## RESULTS

### Early morphine withdrawal increases hedonic feeding

Experiment 1 investigated the effect of morphine withdrawal on hedonic sucrose intake (Fig. 1A and 1B). Prior to drug exposure, sucrose intake increased with sucrose concentration (F_3,33_ = 24.01, < .001) and did not differ across planned drug groups (F < 1; Fig. 1C). The morphine group then received repeated saline and morphine injections paired with distinct unpaired and paired contexts, respectively, whereas the saline group received saline in both contexts (see Fig. 1A and 1B). Morphine initially suppressed locomotor activity in the paired context (Day x Group: F_10,110_ = 3.05, p = .002; Fig. 1D). Rats lost weight during morphine exposure (Day x Group: F_10,110_ = 47.16, p < .001; Fig. 1E), which was exacerbated during early withdrawal (Day x Group: F_1,11_ = 16.41, p = .002; Fig. 1F). Morphine-treated rats also showed increased somatic withdrawal signs (Fig. 1G for details) on WD1 and WD2 (Day x Group: F_1,11_=70.07, p < .001; Fig. 1H). Initial sucrose intake on WD1 was significantly elevated in the morphine group (t_11_ = 3.30, p = .007; Fig. 1I, left). However, this group difference was no longer apparent on WD2, nor was there any apparent influence of drug context (F’s < 1; Fig. 1I, right).

### Motivation and goal-directed control are impaired during early morphine withdrawal

Experiment 2 investigated the effect of morphine withdrawal on instrumental responding for palatable food rewards (Fig. 2A and 2B). Baseline press rates at the end of training did not significantly differ across planned groups (F_1,17_ = 3.26, p = 0.089) or devaluation conditions (F < 1; Fig. 2C). Rats were then treated with morphine and/or saline as in Exp. 1, again leading to locomotor suppression after early morphine injections (Day x Group for paired sessions: F_10,170_ = 3.12, p = .001; Fig. 2D).

Rats were then administered a pair of reward devaluation tests to assess rats’ ability to select instrumental actions in a flexible, goal-directed manner (Fig. 2A). Brief morphine re-exposure was provided between tests to keep the withdrawal interval fixed, such that both tests (one in paired and one in unpaired context) were conducted on WD2. Rats received instrumental retraining on WD1 to assess their motivation to work when both food rewards were highly valued. Retraining press rates were generally depressed in the morphine group (F_1,17_ = 9.78, p = .006; Fig. 2E left), which was was also apparent after normalizing for individual differences in baseline performance (t_17_ = 3.32, p .004; Fig. 2E right).

Prior to testing, rats were fed to satiety on one of the two food rewards (Fig. 2F). The morphine group consumed marginally less food (t_17_ = 1.96, p = .067; Fig. 2G; see Fig. S1 for discussion). The test began with an *Extinction* phase, allowing us to probe rats’ ability to flexibly choose between actions based on *expected outcomes* (Fig. 2H; left). Rats selectively reduced their rate of responding for the devalued reward (Devaluation: F_1,17_ = 18.56, p < .001), an effect that varied with group (Group x Devaluation: F_1,17_ = 4.64, p = .046) but not context (other F’s ≤ 1.37; p’s ≥ .26). While both groups displayed a devaluation effect, this effect was larger (Group x Devaluation: partial η^2^ = .214) for the saline group (F_1,9_ = 14.20, p = .004; partial η^2^ = .612) than for the morphine group (F_1,8_ = 5.82, p = .042; partial η^2^ = .421). Relative to controls, the morphine group showed a significantly lower rate of responding for the nondevalued (F_1,17_ = 5.0, p = .039) but not for the devalued reward (F<1).

In the *Reinforced* test phase (Fig. 2H; right), which allowed us to assess the rats’ ability to use *experienced outcomes* to modify their behavior, overall press rates were significantly lower in the morphine group (F_1,17_ = 6.04, p = .025). The morphine group also showed continued insensitivity to reward devaluation (Devaluation x Group: F_1,17_ = 5.37, p = 0.033; partial η^2^ = .24; devaluation effect in saline group: F_1,9_ = 17.18, p = .003; devaluation effect in morphine group: F_1,8_ = 1.54, p = .25) despite receiving feedback about current reward values (other F’s ≤ 1.40; p’s ≥ .25). Relative to controls, the morphine group responded at a lower rate for the nondevalued (F_1,17_ = 7.03, p = .017) but not the devalued (F<1) reward.

### Motivation and goal-directed control are restored after protracted morphine withdrawal but impaired again after brief morphine re-exposure

Experiment 3 investigated motivation and goal-directed control after protracted morphine withdrawal. Rats in this experiment trained as in Experiment 2. Baseline response rates (Fig. 3B) did not differ across planned drug or devaluation conditions (F’s < 1). Rats then received daily morphine and/or saline treatment as in Experiments 1 and 2, resulting in locomotor suppression following early morphine exposure (Day x Group for paired sessions: F_10,240_ = 17.03, p < .001; Fig. 3C).

After protracted morphine withdrawal (WD19-23), rats received instrumental retraining (see Fig. 3A). Press rates did not significantly vary across groups or planned devaluation conditions (F’s < 1), although the morphine group had marginally elevated responding after normalizing for baseline response rate (t_24_=1.85, p = 0.077; Fig. 3D).

Pre-test food intake did not differ between groups (t_24_ = 1.48, p = .15; Fig. 3E). During the *Extinction test phase*, morphine and saline groups responded at similar levels (F<1; Fig. 3F, left) and selectively withheld the action associated with the devalued reward to a similar degree (F_1,24_ = 36.61, p < .001). Although a Context x Devaluation interaction was detected (F_1,24_ = 4.73, p = .04), this effect did not vary with group (Context x Group x Devaluation: F < 1) and was thus not attributable to morphine expectancy (F’s <1 for all other effects and interactions).

Both groups remained sensitive to devaluation during the *Reinforced test phase* (Devaluation: F_1,24_ = 43.21, p < .001; Devaluation x Group: F_1,24_ = 1.72, p = .20; Fig. 3F, right). Interestingly, the morphine group had generally higher rates of responding (Group: F_1,24_ = 4.66, p = .041), which marginally interacted with test context (F_1,24_ = 3.68, p = .067; other F’s ≤ 1.34; p’s ≥ .26).

Rats were then briefly re-exposed to morphine (and/or saline) before undergoing retraining and devaluation testing in a state of acute withdrawal (see Fig. 3A). During re-exposure, morphine now increased locomotor activity (Group x Day for paired sessions: F_4,88_ = 4.44, p = .003; Fig. 4A). As in Exp. 2, retraining press rates during early morphine withdrawal (WD1) were generally suppressed (Group: F_1,22_ = 5.16; p = .032; other F’s ≤ 1.50; p’s ≥ .23; Fig. 4B).

**Figure 4.**
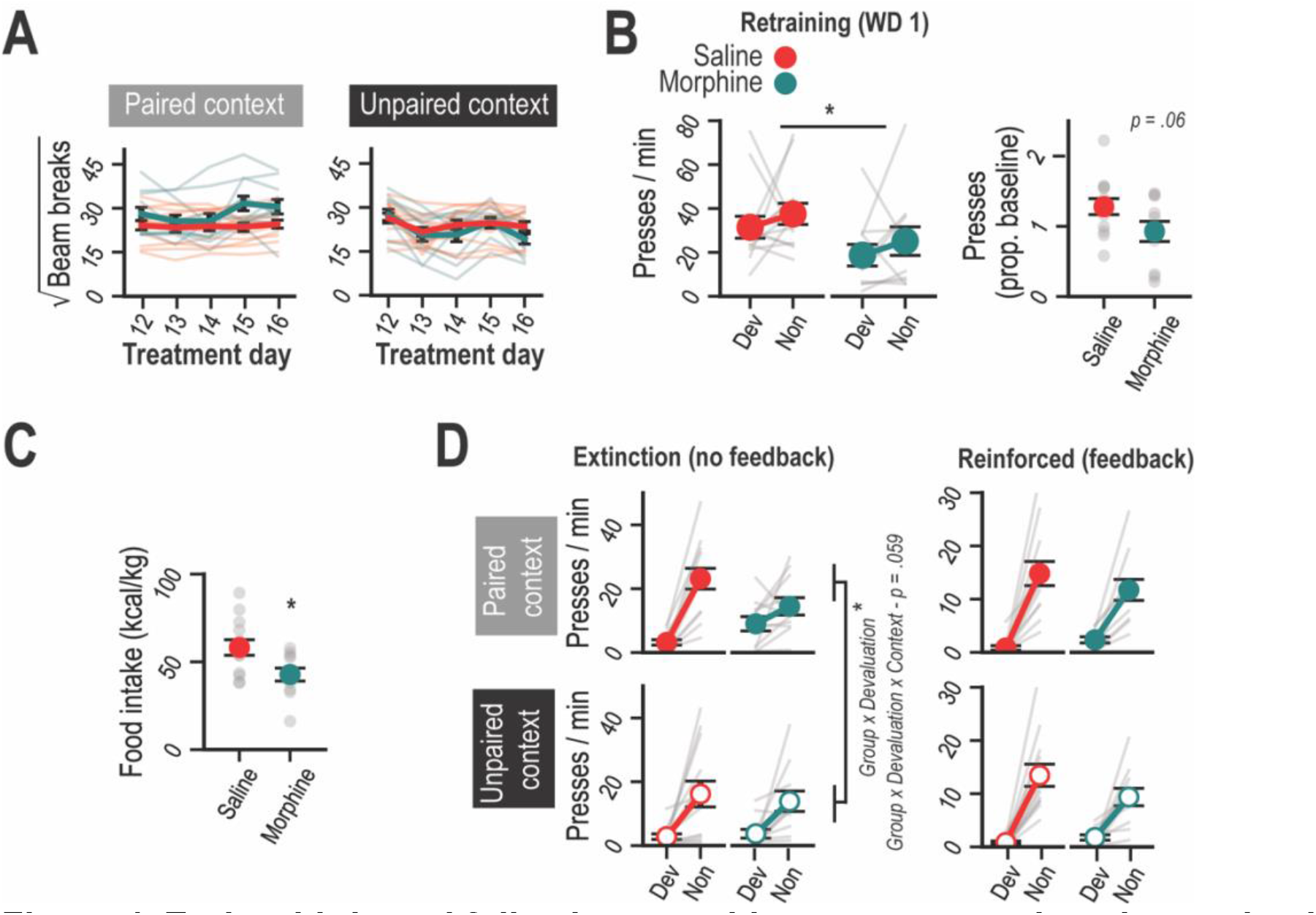
Early withdrawal following morphine re-exposure impairs motivation and goal-directed control in rats. (see Figure 3 for schematic of experimental design). **A**. Locomotor activity during morphine and saline re-exposure. Treatments were given in the original paired or unpaired contexts. Morphine increased locomotor activity in an experience-dependent manner (Day x Group x Context F_4,88_ = 2.71, p = .035), which was more apparent in paired (Day x Group F_4,88_ = 4.44, p = .003) than in unpaired sessions (Day x Group F_4,88_ = 2.52, p = 0.048). **B**. Instrumental performance was depressed in morphine-treated rats during instrumental retraining on WD1 following morphine re-exposure. Left, press rates in rats are separated by group and by planned devaluation conditions. General suppression of lever press rate (Group effect: F_1,22_ = 5.16, p = .032) is not dependent on planned devaluation conditions (Devaluation: F_1,22_ = 1.50, p = .23; Group x Devaluation: F < 1). Right, press rates were averaged across actions and plotted as a proportion of baseline performance showing a marginal suppression after morphine retreatment (t_22_ = 1.96, p = .06). **C**. Food intake during the prefeeding period (specific-satiety induction) is reduced in the morphine group (t_22_ = 2.61, p = .016). **D**. Morphine-treated rats show a deficit in sensitivity to reward devaluation that was specific to the extinction phase of the test that was conducted in the Paired context (see main text for statistical analysis). Press rates during the test session are separated by devaluation (Dev or Non), group (Morphine or Saline), test phase (Extinction and Reinforced), and context (Paired or Unpaired). Dev = Devalued; Non = Nondevalued. *p < .05.

The morphine group consumed less food during the pre-feeding period (t_22_ = 2.61, p = .016; Fig. 4C; see Fig. S1). During the *Extinction* phase of the devaluation test (Fig. 4D, left), sensitivity to reward devaluation was disrupted in the morphine group (Devaluation x Group: F_1,22_ = 5.20, p = .032). There was also a marginally significant interaction between Devaluation x Group x Context (F_1,22_ = 3.97, p = .059). Both groups responded similarly in the unpaired context (Devaluation: F_1,22_ = 19.45, p < .001; Devaluation x Group: F < 1) but were differentially sensitive to devaluation in the paired context (Devaluation x Group: F_1,22_ = 11.83, p = .002). Whereas the saline group displayed a selective devaluation effect in the paired context (F_1,12_ = 50.52, p < .001), the morphine group did not (F_1,10_ = 2.95, p = .12). Rats also showed sensitivity to devaluation during the *Reinforced* test phase (F_1,22_ = 78.18, p < .001) regardless of group or context (other F’s ≤ 1.50; p’s ≥ .23).

Please see Supplementary Information for detailed results.

## DISCUSSION

We investigated how morphine withdrawal impacts processes controlling the pursuit and consumption of palatable food rewards. Rats experiencing acute morphine withdrawal displayed elevated levels of hedonic feeding behavior but showed reduced motivation and impaired goal-directed control over their food-seeking instrumental actions. These deficits were no longer apparent after prolonged morphine withdrawal but were at least partly reinstated following a brief period of morphine re-exposure. We believe these findings have implications for understanding and studying goal-narrowing in opioid addiction.

A defining feature of addiction is that abruptly discontinuing drug use can trigger a so-called *motivational withdrawal syndrome* [6,57,58], which is thought to involve a range of negative emotional changes (dysphoria, irritability and anxiety) and a loss of sensitivity to (anhedonia) and motivation for (amotivation) non-drug rewards. Our findings align with previous reports that motivation for palatable food rewards is attenuated during the early phase of opioid withdrawal [59-61]. Although persistent motivational deficits have also been observed after more protracted periods of opioid withdrawal [21,23,62], other findings suggest that motivational function recovers over time [60,61]. There have also been numerous reports of food-motivated behavior becoming elevated during opioid withdrawal, particularly after long drug-free periods [25,26,28,30,59,60,63,64], which is more compatible with an incentive-sensitization theory of addiction [65,66]. Consistent with this, we found that after prolonged morphine withdrawal rats displayed heightened motivation for food reward during instrumental retraining sessions and during the reinforced phase of the devaluation test. Such findings suggest that opioid withdrawal triggers a biphasic pattern of motivational change in which an initial apathy-like motivational deficit is supplanted by a more persistent state of heightened incentive motivation.

Our findings also address the impact of morphine withdrawal on feeding behavior. We show that total food consumption during 1-h free-feeding sessions (prior to devaluation testing) was reduced during early but not after protracted morphine withdrawal, which is consistent with previous research [31-34]. However, this reduction in feeding does not appear to reflect a deficit in processing the hedonic-emotional properties of palatable food stimuli. We found that rats experiencing early morphine withdrawal showed a significant elevation rather than a reduction in initial, hedonically-driven sucrose intake (prior to satiety). This fits with prior research showing unchanged or heightened hedonic responses to food stimuli [29,35]. Previous research also indicates that opioid withdrawal suppresses consumption of standard maintenance diet and tap water [31,33], and preferentially reduces intake of low-versus high-palatability sucrose solutions [59]. We therefore suggest that hedonic food processing is not impaired and may be upregulated during early morphine withdrawal, even when other factors (e.g., motivational deficit or increased satiety) reduce total consumption. This combination of decreased homeostatic feeding and increased hedonic feeding may explain the generally poor nutrition and increased preference for sugary foods displayed by chronic opioid users [67-69]. Such findings also raise questions about whether opioid withdrawal induces a state of *true* anhedonia (i.e., a decrease in the experience of pleasure when consuming a reward stimulus) or whether other affective, motivational, or even cognitive changes produce symptoms interpreted as anhedonia [70-72].

We also provide critical new information about the impact of morphine withdrawal on goal-directed action selection. Rats in early morphine withdrawal were impaired in flexibly adjusting their choice between actions following reward devaluation, regardless of whether they were tested in a morphine-paired or unpaired context. This impairment was observed both when rats were forced to rely solely on previously learned action-outcome associations to guide their choice (*extinction* test phase) as well as when they were given response-contingent feedback about the consequences of their actions (*reinforced* test phase). This is notable because prior research indicates that healthy, drug-naïve rats will rapidly re-exert goal-directed control and suppress their habitual behavior it is actually produces a devalued reward [73]. Persistent responding for reward despite negative feedback is therefore indicative of a profound loss of goal-directed control and not simply a bias favoring habitual control [45].

Insensitivity to reward devaluation during reinforced testing has been observed in rats with lesions of the dorsomedial striatum [74], a key component of the brain’s goal-directed behavioral control system [75], as well as in rats with a history of repeated cocaine [76] or methamphetamine [77]. However, such impairments are more commonly observed in animals trained on a simple one action-outcome training protocol that promotes habit formation. The more complex two action-outcome training protocol used here is known to prevent habitual control [78-80]. Previous studies using this more complex task have found rats’ capacity to choose between actions based on reward value is not significantly altered by prior cocaine [53] or amphetamine [81] exposure. Given these null results, the loss of goal-directed control during early morphine withdrawal reported here is particularly notable and indicative of a profound impairment.

However, this impairment was not permanent, in that rats tested after an extended withdrawal period displayed normal sensitivity to reward devaluation. This state was also associated with heightened rather than diminished motivation for food reward. Thus, the impairments in motivation and goal-directed control observed during early morphine withdrawal were relatively short-lived and not the result of long-term cognitive dysfunction. Nevertheless, morphine-experienced rats remained vulnerable to future cognitive impairment. Following a protracted drug-free period, brief exposure to morphine reestablished an opioid-dependent state such that acute withdrawal once again impaired motivation and goal-directed control. However, in this case the deficit in devaluation sensitivity was restricted to the extinction test phase, suggesting a more limited loss of goal-directed control than the feedback-insensitive deficit observed after initial withdrawal from chronic morphine. This conclusion is also supported by the context-specificity of the deficit observed after brief morphine re-exposure, which was only apparent when rats were tested in the morphine-paired context. Interestingly, similar context-specific deficits in reward devaluation sensitivity have been observed when rats are tested in the presence of cues that predict other salient stimuli including alcohol [55], methamphetamine [77], or unrestricted access to highly-palatable junk foods [82]. Together, such findings suggest that affectively-charged environmental cues can perturb the cognitive processes responsible for goal-directed action selection. Such effects may be particularly relevant to understanding mechanisms of relapse, which is known to be strongly influenced by drug-related cues [83].

However, there are some important caveats to the current study. First, since we exclusively used adult male rats as subjects, our findings do not address how factors such as sex and age influence the impact of opioid withdrawal on reward pursuit and consumption. Future research should address these questions, particularly given previous reports of sex- [84-86] and/or age-dependent [87,88] effects on other measures of morphine withdrawal. Another consideration when interpreting the current findings is that withdrawal-induced alterations in food intake and motivation may have interfered with our ability to fully and accurately characterize the impact of withdrawal on goal-directed action selection. However, a more in-depth analysis of individual differences in these measures suggests this was not the case [**Figure S1**]. For instance, although morphine-withdrawn rats tended to consume less than control rats during prefeeding sessions, variability in this measure was not associated with sensitivity to reward devaluation during subsequent choice tests in either group. Likewise, the loss of devaluation sensitivity was not likely caused by a simple floor effect, since deficits in choice were not associated with low rates of responding at test.

We suggest that the motivational and cognitive deficits reported here may contribute in important and potentially distinct ways to goal-narrowing in opioid addiction. For instance, a drop in motivation may reduce the amount of time and effort that one is willing to invest when pursuing goals, whereas a cognitive deficit impacting goal-directed control may instead prevent one from selecting and maintaining adaptive goals to pursue. These deficits may also interact in important ways. For instance, rather than inducing a global impairment in goal-directed control, opioid withdrawal may selectively reduce the cognitive resources allocated to pursuing non-drug goals based on their relatively low value versus the more highly valued goal of using opioid drugs. This interpretation is also compatible with a growing body of evidence that putatively compulsive drug-seeking behaviors may actually represent highly motivated and goal-directed rather than habitual actions [89-91]. Future studies will be needed to refine our understanding of how these motivational and cognitive effects of opioid withdrawal interact and how they specifically contribute to the addiction cycle. Ultimately, identifying the unique motivational and cognitive needs of recovering addicts may be critical for developing more effective, patient-focused addiction therapies.

## Acknowledgements

We thank Alyssa Chan, Haena Lim and Kasuni Bodinayake for their assistance with morphine withdrawal scoring and behavioral testing.

## Author contributions

BH and SBO designed and conceptualized the study, performed the data analyses, and wrote the manuscript. SA, CH, BH, and ZS carried out the experiments. All authors reviewed the content and approved the final version prior to submission.

## References

1 Florence C, Luo F, Rice K. The economic burden of opioid use disorder and fatal opioid overdose in the United States, 2017. Drug Alcohol Depend. 2021;218:108350.

2 Strang J, Volkow ND, Degenhardt L, Hickman M, Johnson K, Koob GF, et al. Opioid use disorder. Nat Rev Dis Primers. 2020;6(1):3.

3 American Psychiatric Association. (American Psychiatric Publishing, 2013).

4 Luscher C, Robbins TW, Everitt BJ. The transition to compulsion in addiction. Nat Rev Neurosci. 2020;21(5):247–63.

5 Kalivas PW, Volkow ND. The neural basis of addiction: a pathology of motivation and choice. Am J Psychiatry. 2005;162(8):1403–13.

6 Koob GF. The dark side of emotion: the addiction perspective. Eur J Pharmacol. 2015;753:73–87.

7 Kiluk BD, Yip SW, DeVito EE, Carroll KM, Sofuoglu M. Anhedonia as a key clinical feature in the maintenance and treatment of opioid use disorder. Clin Psychol Sci. 2019;7(6):1190–206.

8 Der-Avakian A, Markou A. The neurobiology of anhedonia and other reward-related deficits. Trends Neurosci. 2012;35(1):68–77.

9 Janiri L, Martinotti G, Dario T, Reina D, Paparello F, Pozzi G, et al. Anhedonia and substance-related symptoms in detoxified substance-dependent subjects: a correlation study. Neuropsychobiology. 2005;52(1):37–44.

10 Martinotti G, Cloninger CR, Janiri L. Temperament and character inventory dimensions and anhedonia in detoxified substance-dependent subjects. Am J Drug Alcohol Abuse. 2008;34(2):177–83.

11 Rapeli P, Kivisaari R, Autti T, Kähkönen S, Puuskari V, Jokela O, et al. Cognitive function during early abstinence from opioid dependence: a comparison to age, gender, and verbal intelligence matched controls. BMC Psychiatry. 2006;6:9.

12 Fu LP, Bi GH, Zou ZT, Wang Y, Ye EM, Ma L, et al. Impaired response inhibition function in abstinent heroin dependents: an fMRI study. Neurosci Lett. 2008;438(3):322–6.

13 Xie C, Shao Y, Fu L, Goveas J, Ye E, Li W, et al. Identification of hyperactive intrinsic amygdala network connectivity associated with impulsivity in abstinent heroin addicts. Behav Brain Res. 2011;216(2):639–46.

14 Zhang M, Liu S, Wang S, Xu Y, Chen L, Shao Z, et al. Reduced thalamic restingstate functional connectivity and impaired cognition in acute abstinent heroin users. Hum Brain Mapp. 2021;42(7):2077–88.

15 Giordano LA, Bickel WK, Loewenstein G, Jacobs EA, Marsch L, Badger GJ. Mild opioid deprivation increases the degree that opioid-dependent outpatients discount delayed heroin and money. Psychopharmacology (Berl). 2002;163(2):174–82.

16 Su B, Li S, Yang L, Zheng M. Reduced response inhibition after exposure to drug-related cues in male heroin abstainers. Psychopharmacology (Berl). 2020;237(4):1055–62.

17 Marissen MA, Franken IH, Waters AJ, Blanken P, van den Brink W, Hendriks VM. Attentional bias predicts heroin relapse following treatment. Addiction. 2006;101(9):1306–12.

18 Zhao Q, Li H, Hu B, Li Y, Gillebert CR, Mantini D, et al. Neural Correlates of Drug-Related Attentional Bias in Heroin Dependence. Front Hum Neurosci. 2017;11:646.

19 Kutlu MG, Gould TJ. Effects of drugs of abuse on hippocampal plasticity and hippocampus-dependent learning and memory: contributions to development and maintenance of addiction. Learn Mem. 2016;23(10):515–33.

20 Welsch L, Bailly J, Darcq E, Kieffer BL. The Negative Affect of Protracted Opioid Abstinence: Progress and Perspectives From Rodent Models. Biol Psychiatry. 2020;87(1):54–63.

21 Harris GC, Aston-Jones G. Altered motivation and learning following opiate withdrawal: evidence for prolonged dysregulation of reward processing. Neuropsychopharmacology. 2003;28(5):865–71.

22 Lieblich I, Yirmiya R, Liebeskind JC. Intake of and preference for sweet solutions are attenuated in morphine-withdrawn rats. Behav Neurosci. 1991;105(6):965–70.

23 Zhang D, Zhou X, Wang X, Xiang X, Chen H, Hao W. Morphine withdrawal decreases responding reinforced by sucrose self-administration in progressive ratio. Addict Biol. 2007;12(2):152–7.

24 Niu H, Zhang G, Li H, Zhang Q, Li T, Ding S, et al. Multi-system state shifts and cognitive deficits induced by chronic morphine during abstinence. Neurosci Lett. 2017;640:144–51.

25 Li Y, Zheng X, Xu N, Zhang Y, Liu Z, Bai Y. The consummatory and motivational behaviors for natural rewards following long-term withdrawal from morphine: no anhedonia but persistent maladaptive behaviors for high-value rewards. Psychopharmacology. 2017;234(8):1277–92.

26 Rouibi K, Contarino A. The corticotropin-releasing factor receptor-2 mediates the motivational effect of opiate withdrawal. Neuropharmacology. 2013;73:41–7.

27 Morisot N, Rouibi K, Contarino A. CRF2 Receptor Deficiency Eliminates the Long-Lasting Vulnerability of Motivational States Induced by Opiate Withdrawal. Neuropsychopharmacology. 2015;40(8):1990–2000.

28 Rouibi K, Contarino A. Increased motivation to eat in opiate-withdrawn mice. Psychopharmacology. 2012;221(4):675–84.

29 Wassum KM, Greenfield VY, Linker KE, Maidment NT, Ostlund SB. Inflated reward value in early opiate withdrawal. Addict Biol. 2016;21(2):221–33.

30 Scheggi S, Braccagni G, De Montis MG, Gambarana C. Heightened reward response is associated with HCN2 overexpression in the ventral tegmental area in morphine-sensitized rats. Behav Pharmacol. 2020;31(2&3):283–92.

31 Houshyar H, Gomez F, Manalo S, Bhargava A, Dallman MF. Intermittent morphine administration induces dependence and is a chronic stressor in rats. Neuropsychopharmacology. 2003;28(11):1960–72.

32 Harvey-Lewis C, Perdrizet J, Franklin KB. The effect of morphine dependence on impulsive choice in rats. Psychopharmacology (Berl). 2012;223(4):477–87.

33 Langerman L, Piscoun B, Bansinath M, Shemesh Y, Turndorf H, Grant GJ. Quantifiable dose-dependent withdrawal after morphine discontinuation in a rat model. Pharmacol Biochem Behav. 2001;68(1):1–6.

34 Gagin R, Cohen E, Shavit Y. Prenatal exposure to morphine alters analgesic responses and preference for sweet solutions in adult rats. Pharmacol Biochem Behav. 1996;55(4):629–34.

35 De Luca MA, Bimpisidis Z, Bassareo V, Di Chiara G. Influence of morphine sensitization on the responsiveness of mesolimbic and mesocortical dopamine transmission to appetitive and aversive gustatory stimuli. Psychopharmacology (Berl). 2011;216(3):345–53.

36 Fatahi Z, Zeinaddini-Meymand A, Karimi-Haghighi S, Moradi M, Khodagholi F, Haghparast A. Naloxone-precipitated withdrawal ameliorates impairment of costbenefit decision making in morphine-treated rats: Involvement of BDNF, p-GSK3-β, and p-CREB in the amygdala. Neurobiology of Learning and Memory. 2020;167:107138.

37 Sweis BM, Redish AD, Thomas MJ. Prolonged abstinence from cocaine or morphine disrupts separable valuations during decision conflict. Nat Commun. 2018;9(1):2521.

38 Withey SL, Doyle RJ, Porter EN, Bergman J, Kangas BD. Discrimination learning in oxycodone-treated nonhuman primates. Drug and Alcohol Dependence. 2020;207:107778.

39 Harvey-Lewis C, Brisebois AD, Yong H, Franklin KBJ. Naloxone-precipitated withdrawal causes an increase in impulsivity in morphine-dependent rats. Behavioural Pharmacology. 2015;26(3):326–29.

40 Harvey-Lewis C, Perdrizet J, Franklin KBJ. The effect of morphine dependence on impulsive choice in rats. Psychopharmacology. 2012;223(4):477–87.

41 Schippers MC, Binnekade R, Schoffelmeer ANM, Pattij T, De Vries TJ. Unidirectional relationship between heroin self-administration and impulsive decision-making in rats. Psychopharmacology. 2012;219(2):443–52.

42 Maguire DR, Gerak LR, France CP. Effect of daily morphine administration and its discontinuation on delay discounting of food in rhesus monkeys. Behavioural pharmacology. 2016;27(2-):155.

43 Balleine BW, Dickinson A. Goal-directed instrumental action: contingency and incentive learning and their cortical substrates. Neuropharmacology. 1998;37(4-5):407–19.

44 Everitt BJ, Robbins TW. Neural systems of reinforcement for drug addiction: from actions to habits to compulsion. Nat Neurosci. 2005;8(11):1481–9.

45 Ostlund SB, Balleine BW. On habits and addiction: an associative analysis of compulsive drug seeking. Drug discovery today: Disease models. 2008;5(4):235–45.

46 Vandaele Y, Janak PH. Defining the place of habit in substance use disorders. Prog Neuropsychopharmacol Biol Psychiatry. 2018;87(Pt A):22–32.

47 Marshall AT, Liu AT, Murphy NP, Maidment NT, Ostlund SB. Sex-specific enhancement of palatability-driven feeding in adolescent rats. PLoS One. 2017;12(7):e0180907.

48 Halbout B, Hutson C, Hua L, Inshishian V, Mahler SV, Ostlund SB. Long-term effects of THC exposure on reward learning and motivated behavior in adolescent and adult male rats. Psychopharmacology (Berl). 2023;240(5):1151–67.

49 Davis JD, Perez MC. Food deprivation- and palatability-induced microstructural changes in ingestive behavior. American Journal of Physiology-Regulatory, Integrative and Comparative Physiology. 1993;264(1):R97–R103.

50 Davis JD, Smith GP. Analysis of lick rate measure the positive and negative feedback effects of carbohydrates on eating. Appetite. 1988;11(3):229–38.

51 Piao C, Liu T, Ma L, Ding X, Wang X, Chen X, et al. Alterations in brain activation in response to prolonged morphine withdrawal-induced behavioral inflexibility in rats. Psychopharmacology (Berl). 2017;234(19):2941–53.

52 Gellert VF, Holtzman SG. Development and maintenance of morphine tolerance and dependence in the rat by scheduled access to morphine drinking solutions. J Pharmacol Exp Ther. 1978;205(3):536–46.

53 Halbout B, Liu AT, Ostlund SB. A Closer Look at the Effects of Repeated Cocaine Exposure on Adaptive Decision-Making under Conditions That Promote Goal-Directed Control. Front Psychiatry. 2016;7:44.

54 Halbout B, Marshall AT, Azimi A, Liljeholm M, Mahler SV, Wassum KM, et al. Mesolimbic dopamine projections mediate cue-motivated reward seeking but not reward retrieval in rats. Elife. 2019;8:e43551.

55 Ostlund SB, Maidment NT, Balleine BW. Alcohol-Paired Contextual Cues Produce an Immediate and Selective Loss of Goal-directed Action in Rats. Front Integr Neurosci. 2010;4.

56 Balleine BW, Dickinson A. Signalling and incentive processes in instrumental reinforcer devaluation. The Quarterly Journal of Experimental Psychology Section B. 1992;45(4):285–301.

57 Koob GF. Neurobiology of Opioid Addiction: Opponent Process, Hyperkatifeia, and Negative Reinforcement. Biol Psychiatry. 2020;87(1):44–53.

58 Koob GF, Le Moal M. Drug abuse: hedonic homeostatic dysregulation. Science. 1997;278(5335):52–8.

59 Bai Y, Li Y, Lv Y, Liu Z, Zheng X. Complex motivated behaviors for natural rewards following a binge-like regimen of morphine administration: mixed phenotypes of anhedonia and craving after short-term withdrawal. Front Behav Neurosci. 2014;8:23.

60 Ford RD, Balster RL. Schedule-controlled behavior in the morphine-dependent rat. Pharmacol Biochem Behav. 1976;4(5):569–73.

61 Babbini M, Gaiardi M, Bartoletti M. Changes in fixed-interval behavior during chronic morphine treatment and morphine abstinence in rats. Psychopharmacologia. 1976;45(3):255–9.

62 Becker JAJ, Kieffer BL, Le Merrer J. Differential behavioral and molecular alterations upon protracted abstinence from cocaine versus morphine, nicotine, THC and alcohol. Addict Biol. 2017;22(5):1205–17.

63 Nocjar C, Panksepp J. Prior morphine experience induces long-term increases in social interest and in appetitive behavior for natural reward. Behav Brain Res. 2007;181(2):191–9.

64 Cooper ZD, Shi YG, Woods JH. Reinforcer-dependent enhancement of operant responding in opioid-withdrawn rats. Psychopharmacology (Berl). 2010;212(3):369–78.

65 Robinson TE, Berridge KC. The neural basis of drug craving: an incentive-sensitization theory of addiction. Brain research reviews. 1993;18(3):247–91.

66 Robinson TE, Berridge KC. Review. The incentive sensitization theory of addiction: some current issues. Philos Trans R Soc Lond B Biol Sci. 2008;363(1507):3137–46.

67 Mysels DJ, Sullivan MA. The relationship between opioid and sugar intake: review of evidence and clinical applications. J Opioid Manag. 2010;6(6):445–52.

68 Morabia A, Fabre J, Chee E, Zeger S, Orsat E, Robert A. Diet and opiate addiction: a quantitative assessment of the diet of non-institutionalized opiate addicts. Br J Addict. 1989;84(2):173–80.

69 Nolan LJ, Scagnelli LM. Preference for Sweet Foods and Higher Body Mass Index in Patients Being Treated in Long-Term Methadone Maintenance. Substance Use & Misuse. 2007;42(10):1555–66.

70 Berridge KC. Evolving Concepts of Emotion and Motivation. Frontiers in Psychology. 2018;9.

71 Olney JJ, Warlow SM, Naffziger EE, Berridge KC. Current perspectives on incentive salience and applications to clinical disorders. Current Opinion in Behavioral Sciences. 2018;22:59–69.

72 Rømer Thomsen K. Measuring anhedonia: impaired ability to pursue, experience, and learn about reward. Frontiers in Psychology. 2015;6.

73 Adams CD. Variations in the sensitivity of instrumental responding to reinforcer devaluation. The Quarterly Journal of Experimental Psychology Section B. 1982;34(2b):77–98.

74 Yin HH, Ostlund SB, Knowlton BJ, Balleine BW. The role of the dorsomedial striatum in instrumental conditioning. European Journal of Neuroscience. 2005;22(2):513–23.

75 Balleine BW, Peak J, Matamales M, Bertran-Gonzalez J, Hart G. The dorsomedial striatum: an optimal cellular environment for encoding and updating goal-directed learning. Current Opinion in Behavioral Sciences. 2021;41:38–44.

76 LeBlanc KH, Maidment NT, Ostlund SB. Repeated cocaine exposure facilitates the expression of incentive motivation and induces habitual control in rats. PLoS One. 2013;8(4):e61355.

77 Furlong TM, Supit AS, Corbit LH, Killcross S, Balleine BW. Pulling habits out of rats: adenosine 2A receptor antagonism in dorsomedial striatum rescues methamphetamine-induced deficits in goal-directed action. Addict Biol. 2017;22(1):172–83.

78 Colwill RM, Rescorla RA. Instrumental responding remains sensitive to reinforcer devaluation after extensive training. Journal of Experimental Psychology: Animal Behavior Processes. 1985;11(4):520.

79 Colwill RM, Triola SM. Instrumental responding remains under the control of the consequent outcome after extended training. Behavioural Processes. 2002;57(1):51–64.

80 Holland PC. Relations between Pavlovian-instrumental transfer and reinforcer devaluation. J Exp Psychol Anim Behav Process. 2004;30(2):104–17.

81 Shiflett MW. The effects of amphetamine exposure on outcome-selective Pavlovian-instrumental transfer in rats. Psychopharmacology (Berl). 2012;223(3):361–70.

82 Kendig MD, Cheung AM, Raymond JS, Corbit LH. Contexts Paired with Junk Food Impair Goal-Directed Behavior in Rats: Implications for Decision Making in Obesogenic Environments. Front Behav Neurosci. 2016;10:216.

83 Vafaie N, Kober H. Association of Drug Cues and Craving With Drug Use and Relapse: A Systematic Review and Meta-analysis. Jama Psychiat. 2022;79(7):641–50.

84 Bravo IM, Luster BR, Flanigan ME, Perez PJ, Cogan ES, Schmidt KT, et al. Divergent behavioral responses in protracted opioid withdrawal in male and female C57BL/6J mice. European Journal of Neuroscience. 2020;51(3):742–54.

85 Luster BR, Cogan ES, Schmidt KT, Pati D, Pina MM, Dange K, et al. Inhibitory transmission in the bed nucleus of the stria terminalis in male and female mice following morphine withdrawal. Addiction biology. 2020;25(3):e12748.

86 Papaleo F, Contarino A. Gender-and morphine dose-linked expression of spontaneous somatic opiate withdrawal in mice. Behavioural brain research. 2006;170(1):110–18.

87 Mohammadian J, Miladi-Gorji H. Age- and sex-related changes in the severity of physical and psychological dependence in morphine-dependent rats. Pharmacology Biochemistry and Behavior. 2019;187:172793.

88 Hodgson SR, Hofford RS, Wellman PJ, Eitan S. Different affective response to opioid withdrawal in adolescent and adult mice. Life Sciences. 2009;84(1):52–60.

89 Heather N. Is the concept of compulsion useful in the explanation or description of addictive behaviour and experience? Addictive behaviors reports. 2017;6:15–38.

90 Hogarth L. Addiction is driven by excessive goal-directed drug choice under negative affect: translational critique of habit and compulsion theory. Neuropsychopharmacology. 2020;45(5):720–35.

91 Vandaele Y, Ahmed SH. Habit, choice, and addiction. Neuropsychopharmacology. 2021;46(4):689–98.

