## Supplementary Information for "Differential effects of acute and prolonged morphine withdrawal on motivational and goal-directed control over reward-seeking behavior"

### SUPPLEMENTARY MATERIAL AND METHODS

#### Apparatus:

Operant behavioral procedures were conducted in identical operant chambers (ENV-007, Med Associates, St Albans, VT, USA), each housed in a sound- and light-attenuated cubicle. A food-delivery port was located at the center of one end-wall of the chamber, 2.5 cm above the stainless-steel grid floor. Separate cups within the food port were used to deliver 0.1-ml infusions of 50% sweetened condensed milk (SCM) solution (Eagle Brand) via a syringe pump located outside of the cubicle or 45-mg grain pellets (BioServ) via an automated pellet dispenser. A photobeam detector positioned across the food-port entrance was used to monitor head entries. Locomotor activity was monitored with four photobeams that were positioned in a horizontal plane ~2 cm above the grid floor. Each chamber was also equipped with two retractable levers positioned to the left and right of the food port. A houselight (3 W, 24 V) at the top of the opposite end-wall provided general illumination and a fan mounted on the cubicle provided ventilation and background noise. All experimental events were controlled and recorded with a 10-msec resolution using MED-PC IV software.

The above description refers to the bare chamber, which served at the training context during instrumental conditioning sessions. During drug administration sessions and behavioral test sessions, we added visual, tactile, and olfactory cues to create two distinctive contexts. For *Context A*, panels with black-and-white vertical stripes were positioned outside the transparent sidewall and door, a PVC perforated sheet covered the grid floor, and a paper towel scented with 0.4 ml of artificial vanilla extract (McCormick and Co. Inc., Baltimore, MD, USA) was placed in the waste pan (below the grid floor). For *Context B*, white panels with black filled circles were placed outside the sidewall and door, the floor was covered with metal mesh sheet, and a paper towel scented with 0.4 ml of artificial lemon extract (McCormick and Co. Inc.) was placed in the waste pan.

Sucrose licking procedures were conducted in a different set of identical operant chambers equipped with a retractable stainless steel 18-gauge gavage needle that served as a delivery spout. The tip of the spout was extended through a 18 x 12 mm oval aperture during sucrose consumption sessions to provided unrestricted access. Sucrose licking responses were continuously recorded during consumption test sessions using a contact lickometer device (ENV-250B, Med Associates, St Albans, VT, USA).

#### Morphine treatment:

Morphine sulfate provided by the NIDA Drug Supply Program was prepared daily in solution with 0.9% sterile saline. For each experiment, the morphine exposure phase was conducted over 11 consecutive days and followed an initial behavioral training phase, as specified below (see Fig. 1A). Rats were given ad libitum access to lab chow and water while in their cages throughout the morphine exposure phase. Morphine exposure groups were given an injection of saline (1 ml/kg) each morning and an injection of morphine each afternoon, separated by approximately 5-6 hours. The dose of morphine was escalated over days with 2 days at 10 mg/kg, 2 days at 20 mg/kg, 1 day at 25 mg/kg, and 6 days at 30 mg/kg, following [1,2]. Saline-only exposure groups were given saline injections in both the morning and afternoon. For both morning and afternoon treatments, rats were injected (s.c.) before being immediately placed in the operant chambers for 30 min. For all groups, morning and afternoon injections were paired with distinct contexts. For the morphine groups, the morning, saline-paired, context was the *Unpaired context* and the afternoon, morphine-paired, context was the *Paired context*; for the saline-only groups, the morning, saline-paired, context was the *Unpaired context* and the afternoon, also saline-paired, context was the *Paired context* (see Fig.1B). Because some rats died because of morphine exposure before termination of their experiment the Ns reported below refer to the number of rats that completed each experiment.

**Experiment 1**

*Sucrose intake training:* Ad-libitum fed rats were trained to lick sucrose from a retractable stainless-steel spout. Training occurred over five daily sessions (30-min each) during which 20% sucrose solution could be continuously self-administered by licking the spout on a FR-1 schedule (15 ul aliquot over 0.25 sec for each lick; as in [3-5]). Additional licks made during sucrose infusions had no consequence (i.e., only licks made when the syringe pump was inactive resulted in a sucrose infusion). Rats were then given four additional 30-min consumption tests with varying sucrose concentrations (0, 2, 10, 20%, one concentration/day, test order randomized across animals). After establishing the concentration-response function, rats were given two additional tests with 2% sucrose to establish a behavioral baseline prior to the morphine exposure phase. Prior to the final baseline sucrose consumption test, rats were habituated to the observational apparatus and procedures that were later used to score signs of morphine withdrawal (see below). For all sucrose consumption sessions, our primary measure was bodyweight-normalized sucrose intake (ml/kg) during the first 3 min of active licking behavior (beginning after the first contact with sucrose), as in our previous publications [3,5], allowing us to selective assay hedonic feeding with minimal influence of satiety [6,7].

*Morphine exposure:* As described in detail above, rats in the morphine group (n = 5, after excluding three rats that died of overdose) received one saline injection and one morphine injection each day for 11 consecutive days. Each treatment was paired with distinct context cues. In contrast, a saline-only group (n = 8) received two saline injections each day, which were also paired with the two distinct contexts.

*Sucrose consumption during withdrawal:* Approximately 24-h following the last injection (WD1), rats underwent morphine withdrawal assessment (see below) and were then immediately placed in behavioral chambers for a sucrose consumption test (30-min), during which they could continuously self-administer 2% sucrose solution by licking a metal drinking spout. The next day (WD2) rats underwent the same procedure (withdrawal assessment followed by sucrose consumption) with the exception that the paired or unpaired contexts were added to the operant chambers where sucrose licking took place. Rats were then re-exposed to morphine (15 mg/kg for one day and 30 mg/kg for the next two days) and/or saline using the same morning/afternoon schedule and context-treatment pairings that were used during initial drug exposure. Rats were then tested 24-h later (WD1) in the bare operant chambers and then on WD2 in the alternative context, so that each rat underwent testing in both the paired and unpaired contexts.

*Morphine withdrawal assessment:* Prior to each sucrose consumption session, on WD1 and WD2, rats were placed in a transparent plastic cylinder (30-cm diameter, 60-cm height) and continuously video recorded over a 30-min observation period. Withdrawal severity scoring was conducted using a rating scale slightly modified from previous studies [8,9], with specific weighting factors shown in Figure1G. Manifestations of withdrawal included body weight loss, jumping or escape attempts, wet-dog shakes, head shakes, tremors, teeth chattering, ptosis, piloerection, and abnormal posture. Behavioral signs of withdrawal were scored by three trained observers blind to the treatment (one observer scored in real time, and two observers scored based on the video recordings). Withdrawal scores after initial exposure and re-exposure to morphine were averaged for each withdrawal interval (i.e., WD1 and WD2).

**Experiment 2**

*Instrumental training:* Rats were food restricted and given two daily sessions of magazine training, during which they received 15 grain pellets and 15 SCM infusions (0.1 ml) delivered in random order using a 90-s random time schedule with the levers retracted. This was followed by 10 days of instrumental training with two distinct action-outcome contingencies (e.g., left press → grain; right press → SCM) (Fig. 2A and B). The left and right lever-press responses were trained in separate sessions each day, at least 60 min apart. Action-outcome contingencies were counterbalanced across subjects. Each session began with the insertion of the appropriate lever and ended after 30 min elapsed or 20 rewards were earned. Lever pressing was reinforced on a fixed ratio-1 (FR-1) schedule for 1 day. Rats were given additional FR-1 sessions, as needed, until they had earned at least 15 rewards with each response within a single session. Rats were then trained with increasingly effortful random ratio (RR) schedules, with 2 days of RR-5, 3 days of RR-10, and 3 days of RR-20.

*Morphine exposure:* After completing instrumental training, rats were given unrestricted access to lab chow in their home cages beginning three days before the morphine exposure phase of the study. As described above, rats were given repeated daily injections of morphine and saline (morphine group; n = 9, after excluding two rats that died) or two daily saline injections (saline-only group; n = 10, after excluding one rat for equipment malfunction) for 11 days, with each treatment paired with a distinct context. Rats were placed back on food restriction beginning on day 10 of the morphine/saline treatment (i.e., 48h prior to instrumental retraining) and remained food restricted for the rest of the experiment.

*Devaluation tests:* Approximately 24 h after the last morphine (or saline) treatment (WD1), rats were given instrumental retraining to reestablish task performance and remind them of the two action-outcome contingencies. These retraining sessions (two sessions per day, one with each action) were identical to the instrumental sessions described above, with the exception that the schedule of reinforcement shifted from FR-1 to RR-20 within the session (three rewards at FR-1, two rewards at RR-5, one rewards at RR-10, and the remainder at RR-20). Retraining sessions lasted 30 min or until 20 rewards were earned. The following day (WD2), rats underwent specific-satiety induced devaluation testing as in previous studies [5,10-13]. Each rat was selectively satiated on grain pellets or SCM (counterbalanced with drug treatment and training contingencies) by providing them with 60 min of unrestricted access to that food in a separate plexiglass cage identical to their home cage. After satiety induction, rats were immediately placed in the instrumental chamber, which was adorned with Context-A cues (either morphine-paired or unpaired, counterbalanced with drug, satiety, and training conditions) (Fig. 2F). After a 10-min context exposure period, both levers were inserted. For the next 5 min

(*Extinction Phase*), rats were able to freely press the left and right lever but received no food reinforcement/feedback. This was immediately followed by a 15-min period (*Reinforced Phase*), during which rats received feedback, as each action was reinforced with its respective outcome (FR-1 for the first five rewards, followed by a RR-20 for the remainder of the session). Rats were then prepared for a second devaluation test in Context-B, which involved re-exposing them to morphine to ensure that they were tested at the same stage of drug withdrawal (as described in Experiment 1). As before, rats received two injections each day with the same context-treatment pairings. A 15 mg/kg dose of morphine was used during the first day of re-exposure, which was increased to a 30 mg/kg dose for the remaining two days of exposure. Rats remained food-restricted during these treatments. Twenty-four hours after the last re-exposure session (WD1), rats were given a session of instrumental retraining (RR-20). On the next day (WD2), rats were given a final devaluation test identical to the first but in the presence of Context-B cues.

**Experiment 3**

*Instrumental training:* Rats were food-restricted and given instrumental training as described in Experiment 2. Ad libitum access to food was then provided following the last training sessions.

*Initial morphine exposure:* As detailed above, rats were given 11 days of separate saline and morphine injections (morphine group: n = 13 after excluding two rats that died) or two separate saline injections (saline-only group: n = 13), which were paired with distinct contexts. Rats remained undisturbed in their home cage for the next 14 days, after which they were placed on food restriction (i.e., 5 days before instrumental retraining).

*Devaluation tests:* During withdrawal days 19-21, rats received instrumental retraining sessions as described in Experiment 2 (see Fig.3A). Pressing was reinforced with the modified progressive ratio (FR1 to RR-20) on the first day and a RR-20 schedule on the next two days. On withdrawal day 22 (WD22), rats were given a devaluation test in Context-A as described in Experiment 2. Rats then received instrumental retraining sessions (FR1 to RR-20 for one day, and RR-20 for two days) before undergoing a second devaluation test on WD26 in the presence of Context-B.

*Morphine re-exposure and devaluation tests:* After characterizing instrumental performance after a prolonged period of morphine withdrawal, we examined how acute withdrawal following a brief period of morphine re-exposure would affect instrumental performance. Thus, rats were given 3 additional days of treatment wherein morphine and saline (or saline only) were once again paired with their respective contexts, using a 15 mg/kg dose of morphine on the first day followed by 2 days of 30 mg/kg morphine. Rats then received one day of instrumental retraining (FR1 to RR-20) on WD1, as in Experiment 2, followed by a devaluation test in Context-A on WD2. Rats were then re-exposed to morphine (30 mg/kg) and saline (or saline only) for two days and given one day of instrumental retraining (FR1 to RR-20) on WD1 before undergoing a final devaluation test in Context-B on WD2. Note that the final group sizes for this test were n =13 for the saline-only group and n = 11 for the morphine group.

**Data analysis:**

Data were analyzed using mixed ANOVAs or unpaired tests, as appropriate, in SPSS(v29). Significance was set at $p < 0.05$. Significant interactions were followed by an analysis of lower-order interactions or simple effects, as appropriate, to identify contributing factors. For sucrose consumption tests, we analyzed bodyweight normalized intake (ml/kg) using ANOVAs with the between-subjects factor Group (morphine vs. saline) and within-subjects factors Concentration (0%, 2%, 10%, 20%) or Context (Paired and Unpaired), as appropriate. For morphine/saline exposure sessions, locomotor activity (total photobeam breaks, square root transformed) was analyzed using Day x Group x Context ANOVAs. Bodyweight (g) was analyzed using a two-way ANOVA with Group and Day as factors. To further expose withdrawal dependent changes in bodyweight, we analyzed weight change as a percent difference from last treatment day using a Group x Day ANOVA. Withdrawal symptoms (composite weighted score) were analyzed with a Group x Day ANOVA. Instrumental response rates (presses per minute) were analyzed using an ANOVA with Group, Devaluation, and Context as factors, as appropriate. To target the effect of withdrawal on instrumental performance, we computed the rate or lever pressing during the most recent (i.e., post-drug exposure) retraining sessions as a proportion of baseline (i.e., pre-drug) press rates during the last three days of instrumental training. For Experiment 3, press rates during the two instrumental training days during late withdrawal were used as baseline to assess the effect of early withdrawal following morphine re-exposure. For Supplementary Figure 1: Unpaired t-tests were used to assess the effect of Drug group on this measure. Consumption of SCM solution and grain pellets during specific-satiety was analyzed as kcal/kg (calorie content- and bodyweight-normalized) using an unpaired test. We assessed overall and group-specific correlations (Pearson, two-tailed) between this consumption measure and sensitivity to devaluation (presses for devalued reward/(total presses for both devalued and nondevalued rewards)). A similar analysis was also performed to assess correlations between devaluation sensitivity and total response rate during devaluation tests. See the Results section for further details.

### EXTENDED RESULTS

#### Experiment 1: Characterizing signs of early morphine withdrawal and its impact on hedonic feeding

*Pretesting:* Rats were trained to voluntarily consume a palatable sucrose solution from a pump-fed drinking spout. The initial sucrose intake during such sessions provides a selective measure of hedonic control over feeding behavior [6,7]. We validated this measure by determining its sensitivity to sucrose concentration. As can be seen in Figure 1C, initial intake increased with sucrose concentration (Concentration: $F_{3,33} = 24.01$, $p < .001$) and did not differ across planned drug treatment groups (F's < 1).

*Drug treatment and effects on locomotor activity:* Rats then received 11 days of treatment with saline in the morning and either morphine or saline in the afternoon (see Fig. 1A for design details). Distinct sets of contextual cues were associated with morning (unpaired context) and afternoon (paired context) treatments (Fig. 1B). Locomotor activity during these sessions was altered by morphine in an experience-dependent manner (Day x Context x Group: $F_{1,110} = 3.55$, $p < .001$) (Fig. 1D). Activity was suppressed after morphine treatment during initial paired context sessions (Day x Group: $F_{10,110} = 3.05$, $p = .002$). In contrast, both groups showed a similar decline in activity over days in the unpaired context (Group: $F_{1,11} = 2.26$, $p = .16$; Day: $F_{10,110} = 122.43$, $p < .001$; Day x Group: $F_{10,110} < 1$).

*Bodyweight:* As can be seen in Figure 1E, rats treated with morphine gradually lost weight during the drug treatment phase (Group x Day (1-11): $F_{10,110} = 47.16$, $p < .001$), which was further exacerbated during the early withdrawal period (% of last treatment day for WD1 and WD2; Group x Day (2): $F_{1,11} = 16.41$, $p = .002$) (Fig.1F) but returned to normal by the 26th day of withdrawal (Group: $F_{1,11} = .25$, $p = .63$).

*Withdrawal symptoms:* The severity of somatic withdrawal signs was characterized on WD1 and WD2 using a composite weighted score (Fig. 1G for details). Morphine-exposed rats displayed significantly more signs of withdrawal at both time points (Group: $F_{1,11} = 70.07$, $p < .001$; Day and Day x Group interaction: F's < 1) (Fig. 1H).

*Sucrose intake:* Hedonic feeding (first 3-min of intake for 2% sucrose solution) was initially tested after a 24-h withdrawal period (WD1) in a bare behavioral chamber (i.e., no context cues). Relative to the control group, morphine-treated rats showed a significant elevation in sucrose intake ($t_{,11} = 3.30$, $p = .007$) (Fig. 1I, left panel). Sucrose licking was assessed again after a 48-h withdrawal period (WD2), this time in the presence of morphine-paired or unpaired context cues. No influence of drug treatment, context, or their interaction was detected (Fs < 1; Fig.1I, right panel).

**Experiment 2: Motivation and goal-directed control during early withdrawal from morphine**

*Instrumental Training:* Rats were first trained to perform two lever-press responses for different food rewards (Fig. 2A and B). Figure 2C shows rats' baseline rate of lever pressing during the final 3 days of instrumental training, prior to the initiation of drug treatment. Although planned drug treatment group assignments were adjusted to balance for baseline press rates, two rats died during morphine treatment and one saline rat was excluded for equipment malfunction (final n's: morphine group = 9; saline group = 10), which by chance resulted in a modest but nonsignificant difference in press rate (Group: $F_{1,17} = 3.26$, $p = 0.089$). Baseline press rates were also balanced across actions with respect to the planned devaluation treatment (Devaluation effect and Devaluation x Group interaction: F's < 1).

*Morphine exposure and locomotion:* Following instrumental training, rats went through the same context-specific morphine (or saline-only) treatment regimen used in Experiment 1. Figure 2D shows the experience-dependent effects of morphine on locomotor activity (Day x Context x Group: $F_{10,170} = 4.58$, $p < .001$). For paired context sessions, morphine treated rats displayed a suppression of activity that was less apparent over days (Day x Group: $F_{10,170} = 3.12$, $p = .001$). The groups did not differ and showed similar rates of habituation of locomotor activity during unpaired context sessions (Group: $F_{1,17} = 3.19$, $p = .09$; Day: $F_{10,170} = 20.75$, $p < .001$; Day x Group: $F_{10,170} = 1.024$, $p < .43$).

*Effects of acute morphine withdrawal on instrumental performance:* Rats were administered a pair of reward devaluation tests to assess whether morphine withdrawal and/or morphine-paired context cues impacted their ability to select instrumental actions in a flexible, goal-directed manner (Fig. 2A). Brief re-exposure to morphine was provided between tests to ensure that both tests were conducted at a 48-h drug withdrawal interval (WD2). One test took place in the paired context and other took place in the unpaired context.

On the day between the last drug treatment and devaluation testing (i.e., at 24-h withdrawal, WD1), rats were given instrumental training to remind them of the instrumental contingencies and assess their motivation to work for food reward. Figure 2E (left panel) shows that press rates were generally depressed in the morphine group ($F_{1,17}$ = 9.78, p = .006) but did not differ according to planned devaluation conditions (Devaluation: $F_{1,17}$ = 1.40, p = .25; Devaluation x Group: F<1), suggesting a general decrease in motivation to work for food. This suppression of lever pressing was also apparent when assessing press rates as a proportion of baseline performance in order to control for pre-existing individual differences (Fig. 2E, right panel; $t_{17}$ = 3.32, p = 0.004).

Specific-satiety induced reward devaluation was achieved by prefeeding rats for 1h on one of the two food rewards before each test session (Fig. 2F). Morphine-treated rats consumed marginally less food (kcal/kg) during this prefeeding period ($t_{17}$ = 1.96, p = .067; Fig. 2G; see *Alternative accounts section and Supplementary Fig. 1* for further discussion). Each devaluation test began with an *Extinction* test phase, during which both levers were inserted into the chamber but were not reinforced. This allowed us to probe rats' ability to flexibly choose between lever-press actions based on expected outcomes. These data are shown in Figure 2H (Left panel). Overall press rates did not significantly differ across groups or context conditions (Group: $F_{1,17}$ = 3.48, p = .08; Context: $F_{1,17}$ = 1.69, p = .21; Group x Context: $F_{1,17}$ = 1.00, p = .31). Press rates were significantly lower for the lever that was associated with the now devalued reward (Devaluation: $F_{1,17}$ = 18.56, p < .001), and this effect of devaluation varied with group (Group x Devaluation: $F_{1,17}$ = 4.64, p = .046) but not test context (Context x Devaluation: $F_{1,17}$ = 1.37, p = .26; Context x Group x Devaluation: F < 1). While both groups showed a reduction in responding for the devalued reward, this effect was larger (Group x Devaluation: partial $\eta^2$ = .214) for the saline group ($F_{1,9}$ = 14.20, p = .004; partial $\eta^2$ = .612) than for the morphine group ($F_{1,8}$ = 5.82, p = .042; partial $\eta^2$ = .421). Relative to controls, the morphine group showed a significantly lower rate of responding for the nondevalued ($F_{1,17}$ = 5.0, p = .039) but not for the devalued (F<1) reward.

The test sessions ended with a *Reinforced* phase, which allowed us to assess the rats' ability to use feedback about the consequences of their actions to modify their lever-press performance. Overall press rates were significantly lower in the morphine group (Group: $F_{1,17}$ = 6.04, p = .025) but did not vary with context (Context: F < 1; Context x Group: $F_{1,17}$ = 1.40, p = .25). The morphine group continued to display a weaker reward devaluation effect (Devaluation x Group: $F_{1,17}$ = 5.37, p = 0.033; partial $\eta^2$ = .24; main effect of devaluation: $F_{1,17}$ = 15.30, p < .001; other F's < 1). Whereas the saline group showed a significant reduction in responding for the devalued reward (Devaluation: $F_{1,9}$ = 17.18, p = .003, partial $\eta^2$ = .66), the morphine group responded on both levers to a similar degree (Devaluation: $F_{1,8}$ = 1.54, p = .25, partial $\eta^2$ = .16) even though the reinforcement contingencies were in effect and provided a continuous reminder about current reward values. This impairment in reward devaluation was, once again, associated with a decrease in responding for the valued ($F_{1,17}$ = 7.03, p = .017) but not the devalued (F<1) reward in the morphine group.

**Experiment 3: Motivation and goal-directed control after protracted morphine withdrawal and during early withdrawal following brief morphine re-exposure**

*Instrumental Training:* As in Experiment 2, rats were first trained on two distinct action-outcome contingencies (Fig. 3A). Baseline response rates during the final 3 days of instrumental training (Fig. 3B) did not differ across planned drug treatment groups or devaluation conditions (all F's < 1).

*Morphine exposure and locomotion:* Rats then received daily treatment with saline in the morning and morphine or saline in the afternoon as in Experiments 1 and 2. Figure 3C shows the influence of morphine on locomotor activity, which again varied across treatment days (Day x Context x Group: $F_{10,240}$ = 9.91, p < .001). For paired context sessions, morphine injections initially suppressed locomotor activity (Day x Group: $F_{10,240}$ = 17.03, p < .001). The groups showed a similar decline in activity over days in the unpaired context (Day: $F_{10,240}$ = 43.55, p < .001; Group and Day x Group interactions: F's < 1).

*Instrumental performance following extended withdrawal:* After a period of protracted morphine withdrawal (19 days), rats were retrained with both instrumental action-outcome contingencies (see Fig. 3A). Press rates during this phase (Fig. 3D, left panel) did not significantly vary across groups or planned devaluation conditions (F<1). However, after normalizing for baseline response rates, the morphine group showed a marginally significant elevation in responding (Fig. 3D, right panel; $t_{24}$=1.85, p = 0.077).

As in Experiment 2, rats then underwent reward devaluation testing in the morphine-paired and unpaired context. Pre-test food intake did not differ between groups ($t_{24}$ = 1.48, p = .15) (Fig. 3E). During the *Extinction phase* of the test, the morphine and saline groups responded at similar levels (Group and Context effects and Group x Context interaction: F's < 1) (Fig. 3F, left panels) and displayed a similar suppression of performance on the lever trained with the now devalued reward (Devaluation: $F_{1,24}$ = 36.61, p < .001; Group x Devaluation: F < 1). Although a modest Context x Devaluation interaction was detected ($F_{1,24}$ = 4.73, p = .04), this effect did not vary with drug group (Context x Group x Devaluation: F<1), indicating that it was not attributable to the morphine-context relationship.

Both groups also selectively withheld the action that led to the devalued reward during the *Reinforced phase* of the test (Devaluation: $F_{1,24}$ = 43.21, p < .001; Devaluation x Group interaction: $F_{1,24}$ = 1.72, p = .20), which was also unaffected by test context (Devaluation x Context interaction: F<1; Devaluation x Context x Group interaction: $F_{1,24}$ = 1.34, p = .26) (Fig. 3F, right panels). Interestingly, morphine-treated rats had generally higher rates of responding during this test phase (Group: $F_{1,24}$ = 4.66, p = .041), which marginally interacted with test context (Group x Context interaction: $F_{1,24}$ = 3.68, p = .067; Context effect: F < 1).

*Effect of acute withdrawal after morphine re-exposure on instrumental performance*: Rats were briefly re-exposed to morphine (and/or saline) before undergoing instrumental retraining and devaluation testing in a state of acute withdrawal (see Fig. 3A; final n's: morphine group n = 11, saline = 13). Morphine altered locomotor activity in an experience-dependent manner (Day x Group x Context $F_{4,88}$ = 2.71, p = .035) (Fig. 4A), which was more apparent in paired (Day x Group $F_{4,88}$ = 4.44, p = .003) than in unpaired sessions (Day x Group $F_{4,88}$ = 2.52, p = 0.048). As in Experiment 2, press rates were generally suppressed after morphine withdrawal during subsequent retraining sessions (Fig. 4B; Group effect: $F_{1,22}$ = 5.16, p = .032) but did not differ across planned devaluation conditions (Devaluation: $F_{1,22}$ = 1.50, p = .23; Group x Devaluation: F < 1). A marginally significant suppression in press rate was observed after normalizing for pre-drug response rates ($t_{22}$ = 1.96, p = .06) (Fig. 4B).

Prior to testing, morphine-treated rats consumed significantly less food during the pre-feeding period ($t_{22}$ = 2.61, p = .016; Fig. 4C; see *Alternative accounts section* for further discussion), which is consistent with the effect observed during early withdrawal in Experiment 2. During the Extinction phase of the devaluation test (Fig. 4D, left panels), the groups displayed similar levels of overall responding (F<1). However, sensitivity to reward devaluation (Devaluation: $F_{1,22} = 39.71$, $p < .001$) was significantly disrupted in the morphine group (Devaluation x Group interaction: $F_{1,22} = 5.20$, $p = .032$). There was also a marginally significant interaction between Devaluation x Group x Context ($F_{1,22} = 3.97$, $p = .059$). Further analysis revealed that the groups responded similarly in the unpaired context (Devaluation: $F_{1,22} = 19.45$, $p < .001$; Devaluation x Group interaction: $F < 1$) but significantly differed in their sensitivity to devaluation in the paired context (Devaluation x Group interaction: $F_{1,22} = 11.83$, $p = .002$). Whereas the saline group displayed a selective devaluation effect in the paired context ($F_{1,12} = 50.52$, $p < .001$), the morphine group did not ($F_{1,10} = 2.95$, $p = .12$).

Sensitivity to reward devaluation was rapidly restored in the morphine group during the Reinforced phase of the test (Fig. 4D, right panels). Rats' tendency to withhold responding for the devalued reward (Devaluation effect: $F_{1,22} = 78.18$, $p < .001$) did not vary across groups or contexts (Devaluation x Group and Devaluation x Group x Context interactions: F's < 1; for all other effects and interactions, $p \geq .15$).

### SUPPLEMENTARY RESULTS

#### ***Alternative accounts of reduced sensitivity to reward devaluation during early withdrawal:***

The results of Experiment 2 and 3 demonstrate that early withdrawal from morphine disrupts rats' tendency to flexibly adjust their choice between actions based on specific-satiety induced reward devaluation. However, rats experiencing early withdrawal also tended to exhibit generally lower response rates (reduced motivation), raising the possibility that a floor effect may have interfered with our ability to accurately measure their sensitivity to reward devaluation. Moreover, rats experiencing acute withdrawal also tended to consume less food during prefeeding sessions. Thus, morphine-withdrawn rats may not have been sufficiently satiated to selectively devalue the prefed food reward. It should be noted that certain features of the data seem to be at odds with these alternative accounts. For instance, in Experiment 3, rats undergoing early withdrawal showed a limited deficit in reward devaluation sensitivity that was restricted to the morphine-paired context and the extinction test phase, despite showing more wide-ranging (context-independent) deficits in instrumental performance and food intake. However, to provide a more thorough assessment of these alternative accounts we performed some additional analyses.

To maximize our statistical power, we pooled data from early withdrawal tests in Experiment 2 and 3. We focused our analysis on data from the extinction phase of reward devaluation tests since withdrawal-induced impairments were detected during this period in both experiments. Data from Experiment 2 were averaged across test contexts since rats in this experiment showed a context-independent impairment. Given the context-specific effect observed in Experiment 3, data from this experiment were restricted to the test conducted in the morphine-paired context. From these data, we quantified each rats' tendency to choose the action that had produced the devalued reward as a proportion of both actions (Devaluation score: Dev/(Dev+Non)) and confirmed that morphine-withdrawn rats displayed a significant impairment in devaluation sensitivity on this measure ($t_{41}$ = 3.95, p = .0003) (Supplementary Fig. 1A). We then assessed if this impairment was caused by a floor effect. Although morphine-withdrawn rats tended lower response rates during retraining sessions (Fig. 2E and 4B), their overall rate of responding during devaluation tests (Dev+Non) was not significantly different from controls (unpaired $t_{41}$ = 1.66, p = .10) (Supplementary Fig. 1B). More importantly, inspection of individual differences in these measures reveals that withdrawal-induced impairments in reward devaluation sensitivity were not associated with low rates of responding at test (morphine group: $r_{18}$ = .12, p = .61; saline group: $r_{21}$ = .09, p = .68; All rats: $r_{41}$= -.05, p = .75) (Supplementary Fig. 1D). Thus, there is no indication that a floor effect interfered with assessment of the reward devaluation effect in morphine-withdrawn rats.

Likewise, our analysis of the individual differences indicates that reduced food intake during satiety induction was not responsible for the loss of sensitivity to reward devaluation during early morphine withdrawal. Although morphine-withdrawn rats consumed significantly less food prior to testing (Supplementary Fig. 1C; $t_{41}$ = 3.29, p = .0021), there was no significant relationship in either group between pre-test food intake and sensitivity to reward devaluation (Supplementary Fig. 1E, morphine group: $r_{18}$ = .36, p = .12; saline group: $r_{21}$ = -.27, p = .21; All rats: $r_{41}$= -.19, p = .22). Indeed, careful inspection of these data indicates that withdrawal-induced deficits in reward devaluation sensitivity were if anything associated with high-levels of food consumption equivalent to that seen in the control group (>30 kcal/kg).

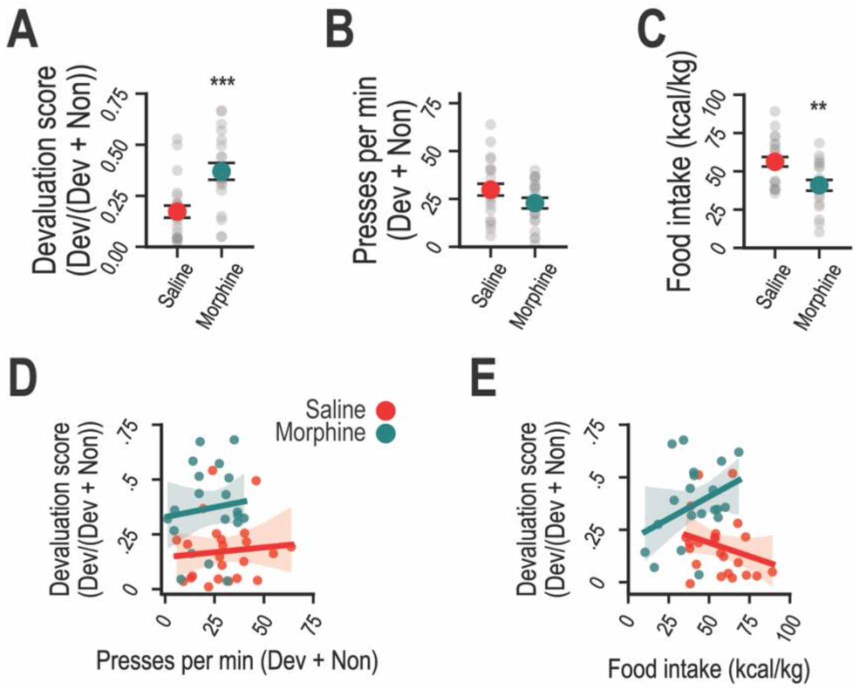


**Supplementary Figure 1. Individual differences in reward devaluation sensitivity do not vary with response rate or prefeeding levels.** Data are combined from early withdrawal tests in Experiments 2 and 3 (see text for details). **A.** Devaluation sensitivity is significantly impaired in the Morphine group. **B.** Overall press rates during the extinction phase of the devaluation test do not significantly differ between Morphine and Saline groups. **C.** Food intake during the prefeeding (specific satiety) period is significantly lower in the Morphine group. **D**. Devaluation sensitivity does not correlate with response rate at test. **E.** Devaluation sensitivity does not correlate with pre-test food intake. Dev = Devalued; Non = Nondevalued. **p < .01. ***p < .001.

### References


1 Wassum KM, Greenfield VY, Linker KE, Maidment NT, Ostlund SB. Inflated reward value in early opiate withdrawal. Addict Biol. 2016;21(2):221-33.

2 Harvey-Lewis C, Perdrizet J, Franklin KB. The effect of morphine dependence on impulsive choice in rats. Psychopharmacology (Berl). 2012;223(4):477-87.

3 Marshall AT, Liu AT, Murphy NP, Maidment NT, Ostlund SB. Sex-specific enhancement of palatability-driven feeding in adolescent rats. PLoS One. 2017;12(7):e0180907.

4 Marshall AT, Halbout B, Liu AT, Ostlund SB. Contributions of Pavlovian incentive motivation to cue-potentiated feeding. Sci Rep. 2018;8(1):2766.

5 Halbout B, Hutson C, Hua L, Inshishian V, Mahler SV, Ostlund SB. Long-term effects of THC exposure on reward learning and motivated behavior in adolescent and adult male rats. Psychopharmacology (Berl). 2023;240(5):1151-67.

6 Davis JD, Perez MC. Food deprivation- and palatability-induced microstructural changes in ingestive behavior. American Journal of Physiology-Regulatory, Integrative and Comparative Physiology. 1993;264(1):R97-R103.

7 Davis JD, Smith GP. Analysis of lick rate measure the positive and negative feedback effects of carbohydrates on eating. Appetite. 1988;11(3):229-38.

8 Piao C, Liu T, Ma L, Ding X, Wang X, Chen X, et al. Alterations in brain activation in response to prolonged morphine withdrawal-induced behavioral inflexibility in rats. Psychopharmacology (Berl). 2017;234(19):2941-53.

9 Gellert VF, Holtzman SG. Development and maintenance of morphine tolerance and dependence in the rat by scheduled access to morphine drinking solutions. J Pharmacol Exp Ther. 1978;205(3):536-46.

10 Halbout B, Liu AT, Ostlund SB. A Closer Look at the Effects of Repeated Cocaine Exposure on Adaptive Decision-Making under Conditions That Promote Goal-Directed Control. Front Psychiatry. 2016;7:44.

11 Halbout B, Marshall AT, Azimi A, Liljeholm M, Mahler SV, Wassum KM, et al. Mesolimbic dopamine projections mediate cue-motivated reward seeking but not reward retrieval in rats. Elife. 2019;8:e43551.

12 Balleine BW, Dickinson A. Signalling and incentive processes in instrumental reinforcer devaluation. The Quarterly Journal of Experimental Psychology Section B. 1992;45(4):285-301.

13 Balleine BW, Dickinson A. Goal-directed instrumental action: contingency and incentive learning and their cortical substrates. Neuropharmacology. 1998;37(4-5):407-19.
